# Reconstruction of consensus tissue-specific metabolic models

**DOI:** 10.1101/327262

**Authors:** Sara Correia, Bruno Costa, Miguel Rocha

## Abstract

Genome-Scale Metabolic Models have shown promising results in biomedical applications, such as understanding cancer metabolism and drug discovery. However, to take full advantage of these models there is the need to address the representation and simulation of the metabolic phenotypes of distinct cell types. With this aim, several algorithms have been recently proposed to reconstruct tissue-specific metabolic models based on available data. Here, the most promising were implemented and used to reconstruct models for two case studies, using omics data from distinct sources. The set of obtained models were compared and analyzed, being shown they are highly variable and that no combination of algorithm and data source can achieve models with acceptable phenotype predictions. We propose an algorithm to achieve a consensus model from the set of models available for a given tissue/cell line, and to improve it given functional data (e.g. known metabolic tasks). The results show that the resulting models are more accurate, both considering the prediction of known metabolic phenotypes and of experimental data not used in the model construction. Two case studies used for model validation consider healthy hepatocytes and a glioblastoma cell line. The open-source implementation of the algorithms is provided, together with the models built, in a software container, allowing full reproducibility, and representing by itself a contribution for the community.

## Author summary

The ability to build models for human cells has the potential to bring powerful tools for biomedical research and drug discovery, since their simulation provides ways to uncover cell phenotypes of both healthy and diseased cells. However, to attain such goals, current models need to improve on their robustness and predictive performance. The growth on the amount and quality of omics data available needs to be put at the service of model reconstruction, dealing with issues such as noise and source heterogeneity. Here, we propose a method to build more reliable genome-scale tissue-specific metabolic models for human cells, using an ensemble-based approach to combine models built from different data sources and using distinct algorithms. We further improve those models using known functional information from the different cell types. This results in models with enhanced capability to predict phenotypes, both in healthy cells (e.g. hepatocyes), and cancer cell lines (e.g. glioblastoma).

## Introduction

Recently, the development of novel techniques for genome sequencing and other high-throughput methods, generating a plethora of “omics” data, such as genomics [1], transcriptomics [2], proteomics [31], metabolomics [3], and fluxomics [4], together with advanced computational methods allowing to mine relevant knowledge from these data sources, boosted the development of Genome-Scale Metabolic Models (GSMMs) of many organisms, including humans. These models are composed by metabolites and reactions that allow the representation of the full set of metabolic processes within a cell, being curated using the knowledge of cellular functions (e.g. from literature). GSMMs, combined with constraint-based approaches, allowed the development of methods to simulate cell behaviour, such as Flux Balance Analysis (FBA) [5]. In turn, these have enabled applications in biotechnology, such as strain optimization [6], but also in the biomedical fields.

Indeed, since the human genome sequencing and annotation [7,8], efforts have been made to reconstruct human genome-scale metabolic models. Until now, five human GSMMs has been proposed [9–13] and have been used to study human physiology and pathology. Just to mention a few illustrative examples, these models have been used to identify drug targets for hypercholesterolemia [9] or markers for inborn errors of metabolism [14], and to try to elucidate the Warburg effect in cancer cell metabolism [15].

However, humans are complex organisms, and the number of distinct cell types is huge, as well as the diversity of their metabolic phenotypes. In spite of the early successes, the full usefulness of human GSMMs lies on the ability to create methods for the phenotype simulation of distinct cell types. During the last decade, a few methods have been proposed towards this aim, integrating generic models with omics data. Here, data are used to generate constraints refining GSMMs, to predict the phenotype of cell types, as it is the case with iMAT [14] and GIMME [16]. However, these only allow the phenotype prediction for conditions for which data was measured, and have revealed less convincing results when tested in prediction benchmarks [17].

An alternative lies in methods that create tissue-specific GSMMs, restricting the metabolic portfolio of the original model (called *template*) to reactions occurring in that cell type, as identified from omics data or biochemical knowledge/ literature. A number of methods have been proposed in the last years to address this task. MBA [18] uses an heuristic approach to prune the original set of reactions, based on evidences from literature and data, and was first used to create an hepatocyte model. Similarly, a variant of GIMME was used to create models for brain cells towards an improved understanding of Alzheimer disease [19]. More recently, the INIT algorithm was applied for the reconstruction of GSMMs for 69 cell types [11], based on mixed integer-linear programming, using proteomics and metabolomics data. The task-driven INIT (tINIT) is an extension of the previous algorithm [20] that allows to define metabolic tasks that the resulting models need to be able to achieve. mCADRE [21] and FASTCORE [22] are based on the ideas of MBA, but reformulate the problem to make the algorithms computationally more efficient, and thus have been used to reconstruct models in a larger scale. A recent method, PRIME [23], uses cell growth measurements, and their correlations with gene expression, to filter the relevant reactions.

These methods have been used in a number of applications, many of which are related to drug discovery research, mainly to identify metabolic targets that can inhibit cancer cell growth. A first study built a generic cancer model [24], using a variant of MBA, proposing a set of generic drug targets. Focused approaches addressed the reconstruction of models for specific cancer types, as renal cancer where a pathway used by cancer cells and not by healthy ones was uncovered, allowing to identify targets for therapy [25].

Still, in spite of these successful applications, no algorithm for tissue-specific reconstruction has shown a consistent ability to generate accurate models, able to provide phenotype predictions according to known metabolic behaviour, taken from literature or available data. Also, the models provided by different methods and data sources have shown a very high level of variability in some preliminary studies [26], which may suggest that an ensemble based approach might lead to better models.

Here, the most promising algorithms (MBA, tINIT, mCADRE and FASTCORE) were used to reconstruct tissue-specific metabolic models for two case studies, hepatocytes and a glioblastoma cell line, resorting to different data sources. The models generated in each case were analyzed and compared in different perspectives.

A new algorithm to combine the set of obtained models was proposed and applied to achieve a consensus metabolic model for each case study. The resulting models were validated using knowledge about known metabolic tasks in liver and omics data not used in the model building process (e.g. metabolomics and growth rates).

## Methods

### Tissue-specific reconstruction methods

The reconstruction of specific metabolic models is a process with four main steps (Fig 1):

1. Collect the data from the repositories and convert the gene identifiers to the notation used in the template metabolic model.
2. Based on the data from the previous step, convert the gene scores to reaction scores. This step can be performed using the gene-protein-reaction (GPR) associations present in the template model, replacing the AND/OR operators by Min/Max functions applied to the respective gene scores.
3. Build the core reaction sets, based on the assumptions described in Table 1. In this step, some additional configurations are required depending on the selected algorithm. In this work, the metabolic tasks used in the tINIT algorithm were retrieved from [20]. Additionally, a set of metabolites were set in mCADRE and tINIT algorithms as described in the original publications [20,21].
4. Finally, one of the selected algorithms is run to reconstruct a tissue-specific metabolic model. Here, the final MBA models were constructed based on 50 intermediate metabolic models. According to [18], a larger number would be desirable, but the time and computational resources needed to generate each model prevented larger numbers of replications.

**Fig 1.**
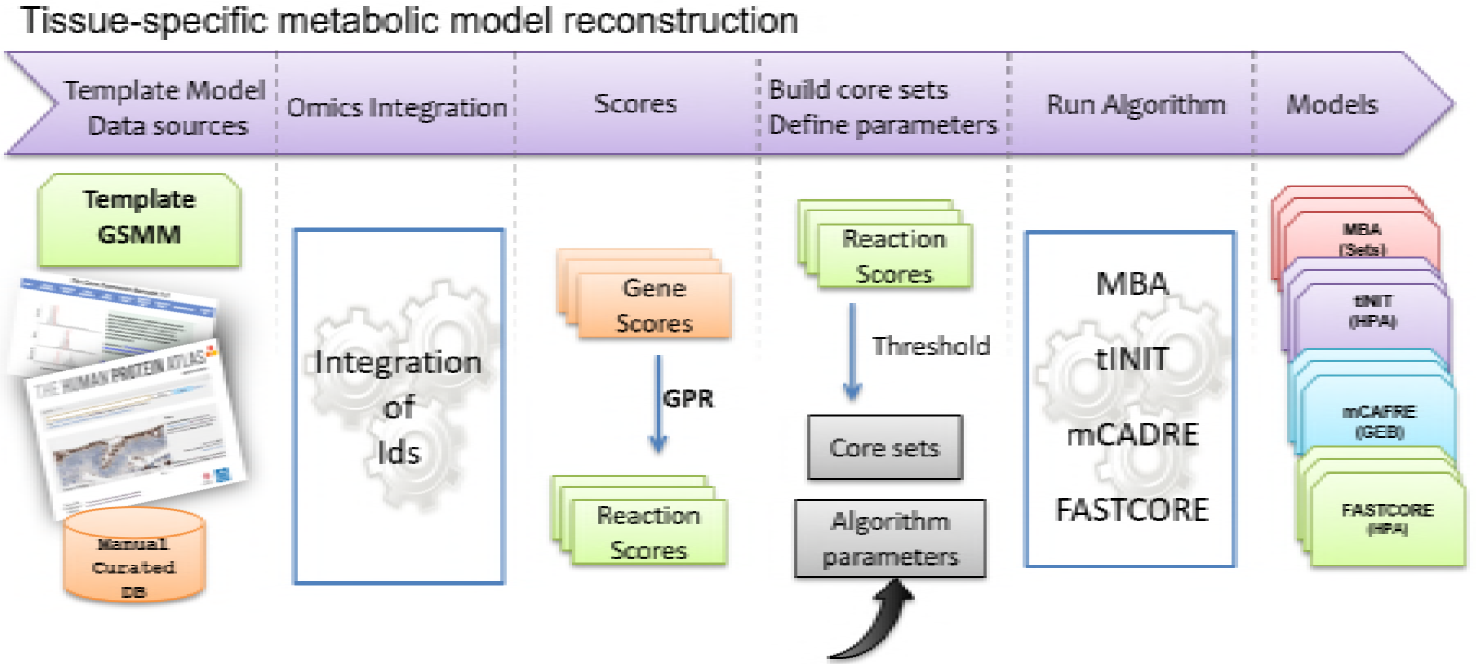
Pipeline for tissue-specific metabolic models reconstruction. The reconstruction of tissue-specific metabolic models has four main steps: collect data; convert scores from gene to reaction level; build the core reactions set and run the selected algorithm.

**Table 1.**
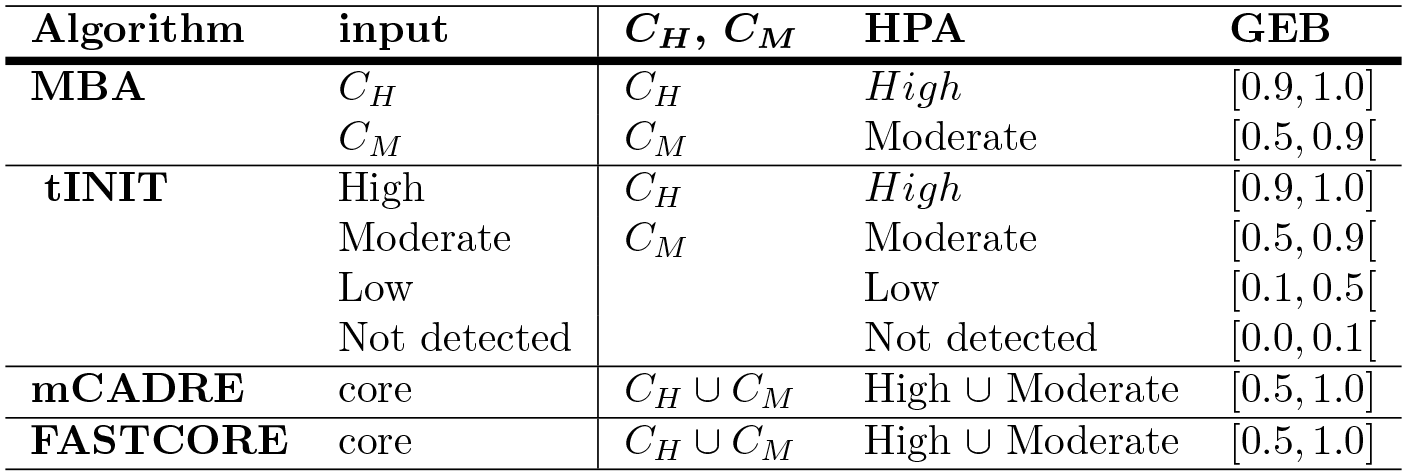
Core sets used as input data.

### MBA

The *Model-Building Algorithm* (MBA) [18] reconstructs a tissue-specific metabolic model from a generic model by integrating a variety of tissue-specific molecular data sources. The first step of this algorithm is to infer, from the tissue-specific data, two sets of reactions denoted as the core reactions (*C*_*H*_) and reactions that have a moderate probability to be carried out in the specific tissue (*C*_*M*_). This division is made according to the accuracy level of the input data. In general, the *C*_*H*_ set includes human-curated tissue-specific pathways and the *C*_*M*_ set includes reactions certified by molecular data. The aim of this method is to find the most parsimonious tissue-specific consistent model, which includes all the tissue-specific high-probability reactions (*C*_*H*_), a maximal number of moderate probability reactions (*C*_*M*_) and a set of additional reactions from the generic model that are required for gap filling, using a greedy heuristic search that is based on iteratively pruning reactions from the generic model.

### tINIT

The *task-driven Integrative Network Inference for Tissues* (tINIT) [20] reconstructs tissue-specific metabolic models based on protein evidence from Human Protein Atlas (HPA) and a set of metabolic tasks that the final context-specific model must perform. The HPA data is the main source of evidence for assessing the presence or absence of metabolic enzymes in each of the human cell types. Moreover, other data sources as tissue specific gene expression and metabolomics data from Human Metabolome Database (HMDB) can also be used. The tasks used by the algorithm aim to test the production or uptake of external metabolites, but also the activation of pathways that occur in a specific tissue. During the validation process for the tasks in the template model, a set of required reactions are found, and constraints to ensure the flux through these reactions are added to the formulation.

### mCADRE

The *Metabolic Context specificity Assessed by Deterministic Reaction Evaluation* (mCADRE) [21] method is able to infer a tissue-specific network based on gene expression data, network topology and reaction confidence levels. Based on the expression score, the reactions of the global model, used as template, are ranked and separated in two sets: core and non-core. All reactions with expression-based scores higher than a threshold value are included in the core set, while the remaining reactions make the non-core set. In this method, the expression scores do not represent the expression levels, but rather the frequency of expressed states over several transcript profiles. Hence, it is necessary to initially binarize the expression data. Thus, it is possible to use data retrieved from the Gene Expression Barcode (GEB) project that already contains binary information on which genes are present or not in a specific tissue/ cell type.

Reactions from the non-core set are ranked according to the expression scores, connectivity-based scores and confidence level-based scores. Then, sequentially, each reaction is removed and the consistency of the model is tested. The elimination only occurs if the reaction does not prevent the production of a key metabolite, i.e. metabolites that have evidence to be produced in the context-specific model reconstruction, and the core consistency is preserved.

## FASTCORE

FASTCORE [22], is a generic algorithm for context-specific metabolic models reconstruction that takes as input a core set of reactions and a generic metabolic model. Firstly, it converts the initial model to a consistent model, i.e. only reactions that can carry flux in at least one feasible flux distribution are preserved. This can be done using existing approaches such as Flux Variability Analysis (FVA) or a new one proposed together with this algorithm for fast consistency check (FASTCC) of a network. Next, it searches for a sub-network from the generic model that contains all reactions present in the core set and a minimal set of additional reactions, necessary to guarantee the consistency of the final model. Some advantages of this algorithm are that it can be applied to integrate different types of “omics” data through the core set compilation by the user, and there is no need to define parameters except the flux threshold *ϵ*, which is used to guarantee the required minimum flux.

### Omics data sources

Transcriptomics are, certainly, the most widely available type of omics data. Using DNA microarrays or RNA-sequencing allows the quantification of gene expression levels in different conditions [27,28]. However, mRNA molecules are not always translated into proteins [29], and therefore the amount of protein produced depends on the gene expression and the current state of the cell. Thus, the knowledge about the amounts of proteins in the cell, provided by proteomics data [30], is of foremost relevance. These data can confirm the presence of proteins and quantify the amount of proteins within a cell.

The input data used by the reconstruction algorithms present in our framework were retrieved from the Human Protein Atlas (HPA) [31] and Gene Expression Barcode (GEB) [32] databases. The information collected from HPA is scored as “supportive” or “uncertain”, depending on the similarity in immunostaining patterns and consistency with protein/gene characterization data. For the present study, only the data classified as “supportive” was used. Moreover, after the conversion of Ensemble gene identifiers to gene symbols, duplicated genes with different evidence levels were removed.

Regarding the GEB data, the conversion to gene expression levels was done considering the average level of probe sets for each gene. The mapping between probe sets and gene symbols was performed using the library “hgu133plus2.db” from Bioconductor. Besides these two data sources, the reaction sets described in [18] were used in the reconstruction of the hepatocytes metabolic model, one of the case studies in this work.

The context-specific reconstruction methods use different formats of input data. Specifically, MBA uses two sets of reactions, where each reaction has *High* or *Moderate* probability to be in the final model, while mCADRE and FASTCORE expect only one set of reactions as input. In tINIT, each reaction from the template metabolic model must have a score value of 20, 15, 10, −8 representing the *High, Moderate, Low* or *Not detected* evidence of protein expression levels, respectively. A default value of −2 is used for reactions without information in the input data.

This diversity in the input formats leads to the need for data transformation, to allow their use in different methods. The continuous data from GEB was classified as *High, Moderate* and *Low*, if the gene expression evidence on that tissue is greater than 0.9, between 0.5 and 0.9, and between 0.1 and 0.5, respectively. The genes with expression evidence below 0.1 were considered not expressed. The core reaction sets used in mCADRE and FASTCORE methods were built considering the union of “High” and “Moderate” gene evidences from HPA, or gene expression evidence greater or equal to 0.5 from GEB, through the GPR association present in the template model.

Table 1 summarizes the assumptions used to create the input data sets for each algorithm. Applying these transformation rules is possible to adapt different input data sources, such as HPA, GEB and *C*_*H*_ and *C*_*M*_ sets, for all methods.

This table summarizes the assumptions and thresholds used to create the sets used as inputs by the different algorithms.

### Consensus algorithm

A consensus metabolic model can be reconstructed based on models obtained by the combination of different algorithms and data sources. The main idea is to build a model starting with the reactions present in most of the models, and iteratively append a set of reactions to the final model so that it will be able to perform all the metabolic tasks given as input to the algorithm.

Assuming a set of *N* algorithms is run with a set of *M* different input data configurations, we have *N* × *M* possible models. Based on these models, the proposed consensus algorithm consists on four main steps:

1. Build *n* = *N* × *M* models (*partial models*), where the *i*^*th*^ model (*i* ∈ {1, 2, …*n*}), hereafter designed as *pModel*_*i*_, contains the reactions present in at least *i* of the original models. Therefore, *pModel*_1_ contains the union of all reactions from the original models, while *pModel*_*n*_ contains only the reactions present in their intersection. The template model with all potential reactions is considered as *pModel*_0_.
2. Run the validation tasks process for each *pModel* and choose a value of *j*, where *pModel*_*j*_ is the smallest model with an acceptable number of valid tasks (user defined threshold).
3. Calculate the reactions and valid task sets that differ between two neighboring models (i.e. models with indexes *i* and *i* + 1). As a result, two lists are created, the lost reaction set (LRS) and the lost metabolic task set (LMT). Each LMT set, *LMT*_*i*_ represents the set of tasks satisfied by *pModel*_*i*_ but not satisfied by *pModel*_*i*+1_ Similarly, each LRS set, *LRS*_*i*_, represents the reactions present in *pModel*_*i*_ and not in *pModel*_*i*+1_
4. Run Algorithm 1 to generate the final model by removing unnecessary reactions to perform the set of tasks. The algorithm starts with *pModel*_*j*_, and taking into consideration the *LRS*_*i*_ and *LMT*_*i*_, finds the reactions of *LRS*_*i*_ that are not required to perform the tasks in *LMT*_*i*_. These reactions compose the *toDel*_*i*_ set. At the end of each iteration, the reactions that do not have an influence in the loss of metabolic tasks, between two partial models (*toDel*_*i*_), are appended to the *LRS_*i*−1_* in the next iteration. The process ends with the processing of *pModel*_0_, in this case the full template model.

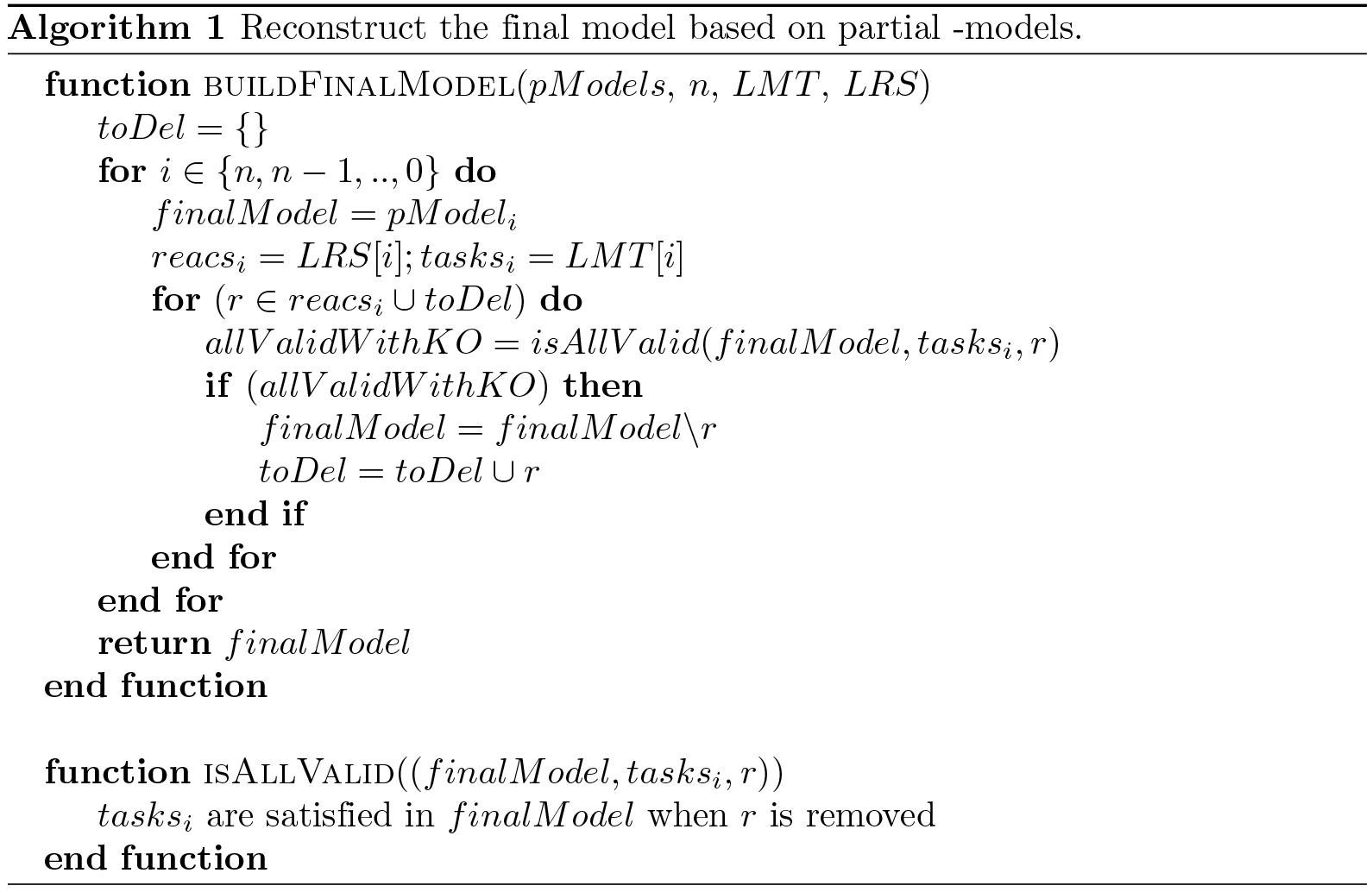

The steps in the reconstruction of the consensus metabolic model algorithm are depicted in Fig 2.

**Fig 2.**
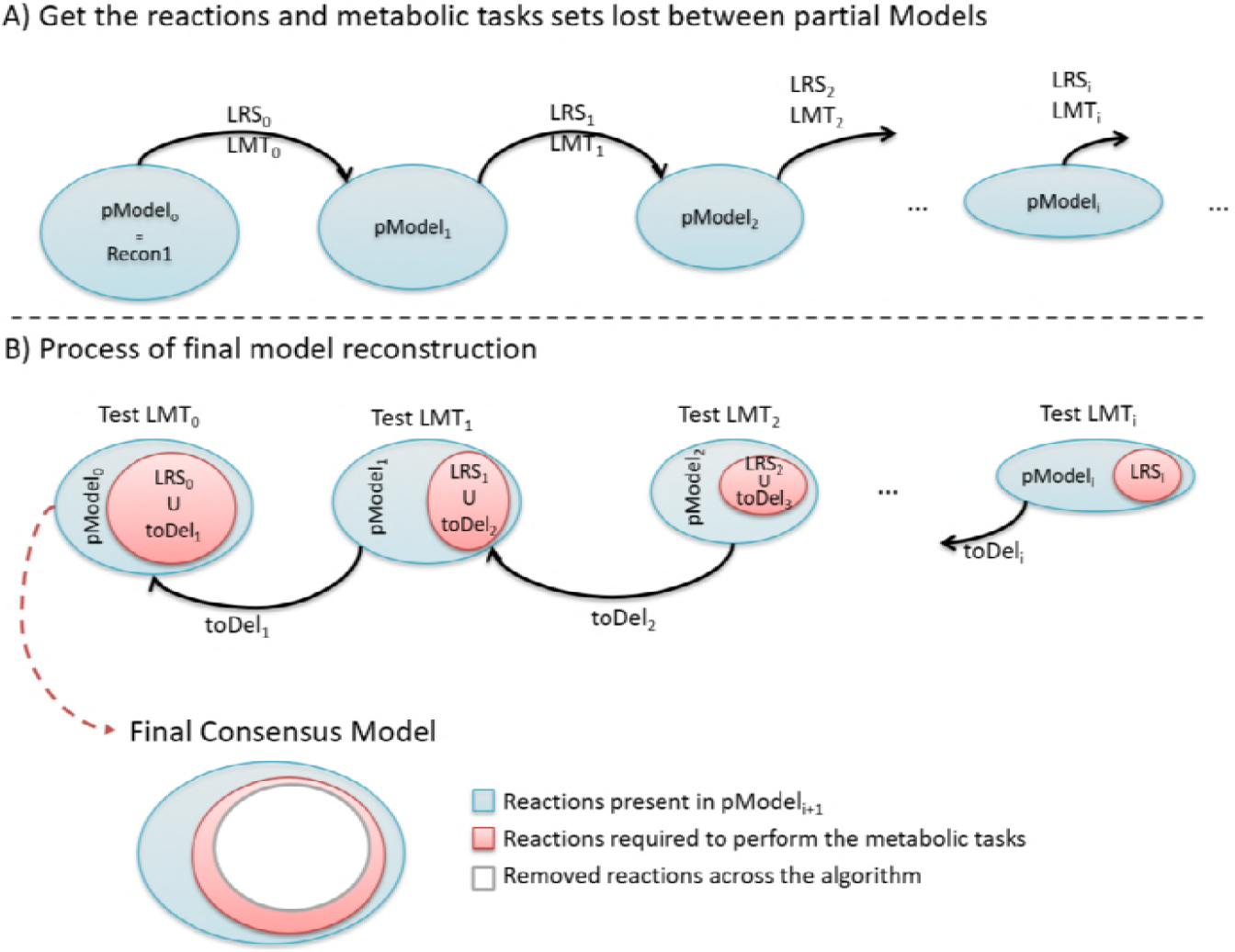
Steps to reconstruct the consensus metabolic model. A) Build the lost reaction set (LRS) and the lost metabolic task set (LMT) between each pair of partial models. B) For each *pModel*_*i*_l the process finds the reactions from *LRS*_*i*_ that can be removed from *pModel*_*i*_ without affecting the metabolic tasks present in (*LMT*_*i*_). The set of reactions that can be removed will be added to the LRS set in the next iteration.

The consensus algorithm here proposed, together with the set of previous algorithms described above, were implemented in the Java language by the authors in an integrated platform to enable comparison of their results. The code is available in GitHub repository (https://github.com/saragcorreia/consensus_tsmm). Moreover, a Docker image are available in the repository Docker Hub (saracorreia/consensus_tsmm/), with all source code, scripts, data and models, for each of the case studies, allowing to fully reproduce the results of this study. These resources represent, by themselves, a contribution of this work, as they make available in a single platform a set of algorithms to reconstruct tissue-specific models that were previously hard to compare since some were available in different platforms (some in commercial platforms as Matlab), while others are not available in any form.

## Results/Discussion

### Case studies

In this study, *Recon 1* was used as template model. The main reasons for this choice are related to the size of the model, being the time consumed to generate the tissue-specific models much lower than using other metabolic models as *Recon 2*. Also, for the glioblastoma case study, this increased the possibilities to compare the results with already published metabolic models [21,23].

About 78% of the liver tissue is formed by hepatocyte cells that are the principal site of the metabolic conversions underlying the diverse physiological functions of the liver [33]. A consensus hepatocyte metabolic model was here reconstructed using different omics data sources and algorithms, also evaluating the effects that each of those data types has in the resulting tissue-specific model.

The hepatocytes metabolic models were generated using *Recon 1* and combining different data sources (GEB, HPA and the *C*_*H*_ and *C*_*M*_ sets from [18]). The four algorithms described above (MBA, tINIT, mCADRE and FASTCORE) were used, leading to the creation of twelve distinct metabolic models were obtained.

Following the same approach, eight metabolic models were reconstructed for U-251 glioblastoma cell lines, considering HPA and GEB as data sources and the same four algorithms previously mentioned. Glioblastoma (GBM), also known as astrocytoma grade IV, is the most common and aggressive type of brain cancer in adults [34]. Despite the advances in the study of this type of cancer, it remains largely incurable.

The consensus algorithm described in the previous section was applied to reconstruct consensus metabolic models for both hepatocytes and U-251 glioblastoma cell lines. In each case, this algorithm took as input the models created using the different algorithms and data sources, as described above.

The consensus models generated in this work are provided in the Systems Biology Markup Language (SBML) format as supporting information.

### Data sources and template model

We start by analysing the data sources contents and their overlap with the metabolic model used as a template. The data retrieved from HPA has evidence for 16324 genes in hepatocytes. On the other hand, the GEB transcriptome (HGU133plus2_cells_v3) has information for 20149 genes, of which 5772 have evidence of being expressed in hepatocytes [32]. Together, these two data sources have information for 21921 genes, but only 14552 are present in both. The gene expression levels present in the omics data sources were converted to reaction evidence levels through the gene-protein-reactions (GPR) rules present in the metabolic model.

Considering these data sources and *Recon 1* as the template model, 1903 reactions show evidence that supports their inclusion in the hepatocytes metabolic model, but only 386 are supported by all sources (Fig 3A; upper row). The numbers are further dramatically reduced if we consider only moderate or high levels of evidence (Fig 3 B and C;upper row).

**Fig 3.**
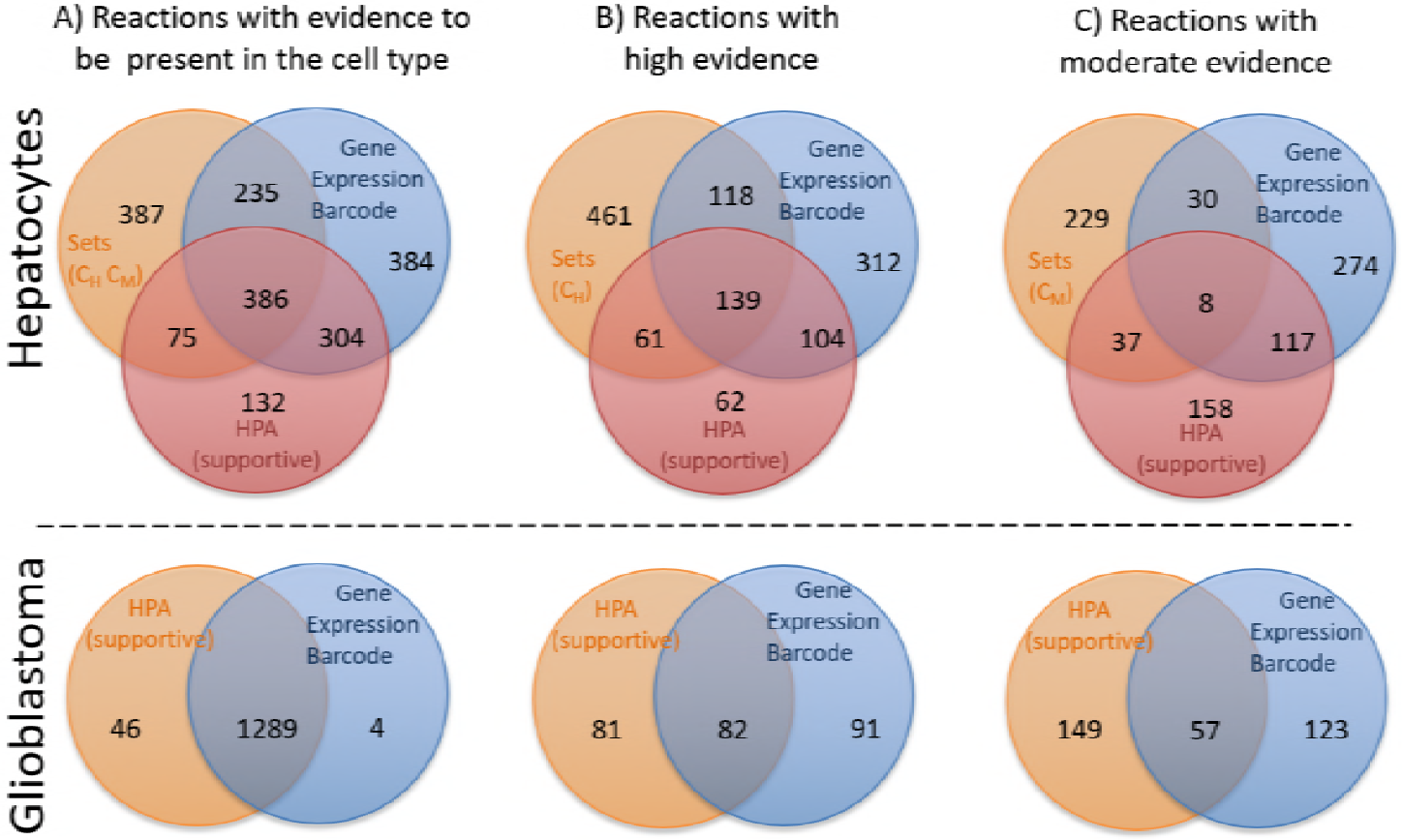
Overlap of reaction evidence levels for the different input data sources (*C*_*H*_ and *C*_*M*_ for hepatocytes, GEB and HPA for both case studies). A) Reactions with evidence that support their inclusion in the tissue-specific metabolic models. B) Number of reactions that have a high level of evidence of expression for each data source. C) Number of reactions that have a moderate evidence of expression for each data source.

Regarding the U-251 cell line, the HPA and GEB databases have information for 1335 and 1293 genes, respectively, from the 1905 genes present in the Recon1 model. Analyzing the reaction evidence levels in this case, the overlap of reactions that support the inclusion in the tissue-specific model is around of 97% and 99% for HPA and GEB, respectively (Fig 3A; lower row). However, if we take into account the expression evidence levels, the number of reactions with the same evidence level is surprisingly low (Fig 3 B and C; lower row). The lower overlap in the high and moderate sets of reactions considering the cutoffs of “High”/ 0.9 and “Medium” / 0.5 from data retrieved from HPA/GEB can have a significant impact in the resulting models, independently of the used algorithm.

### Hepatocytes tissue-specific metabolic models

For the hepatocytes, the twelve metabolic models built in this work have between 1178 and 2139 reactions, as shown in Table 2. A more detailed comparison between the models reconstructed using the same algorithm or the same data source is available in Fig 4, A and B respectively, showing the number of shared reactions in these sets of models.

**Table 2.** Number of reactions for all hepatocytes’ metabolic models.

| | $C_H$ and $C_M$ | HPA | GEB |
| --- | --- | --- | --- |
| MBA | 1748 | 1246 | 1577 |
| tINIT | 1750 | 11837 | 2139 |
| mCADRE | 1760 | 1178 | 1511 |
| FASTCORE | 1817 | 1220 | 1542 |

**Fig 4.**
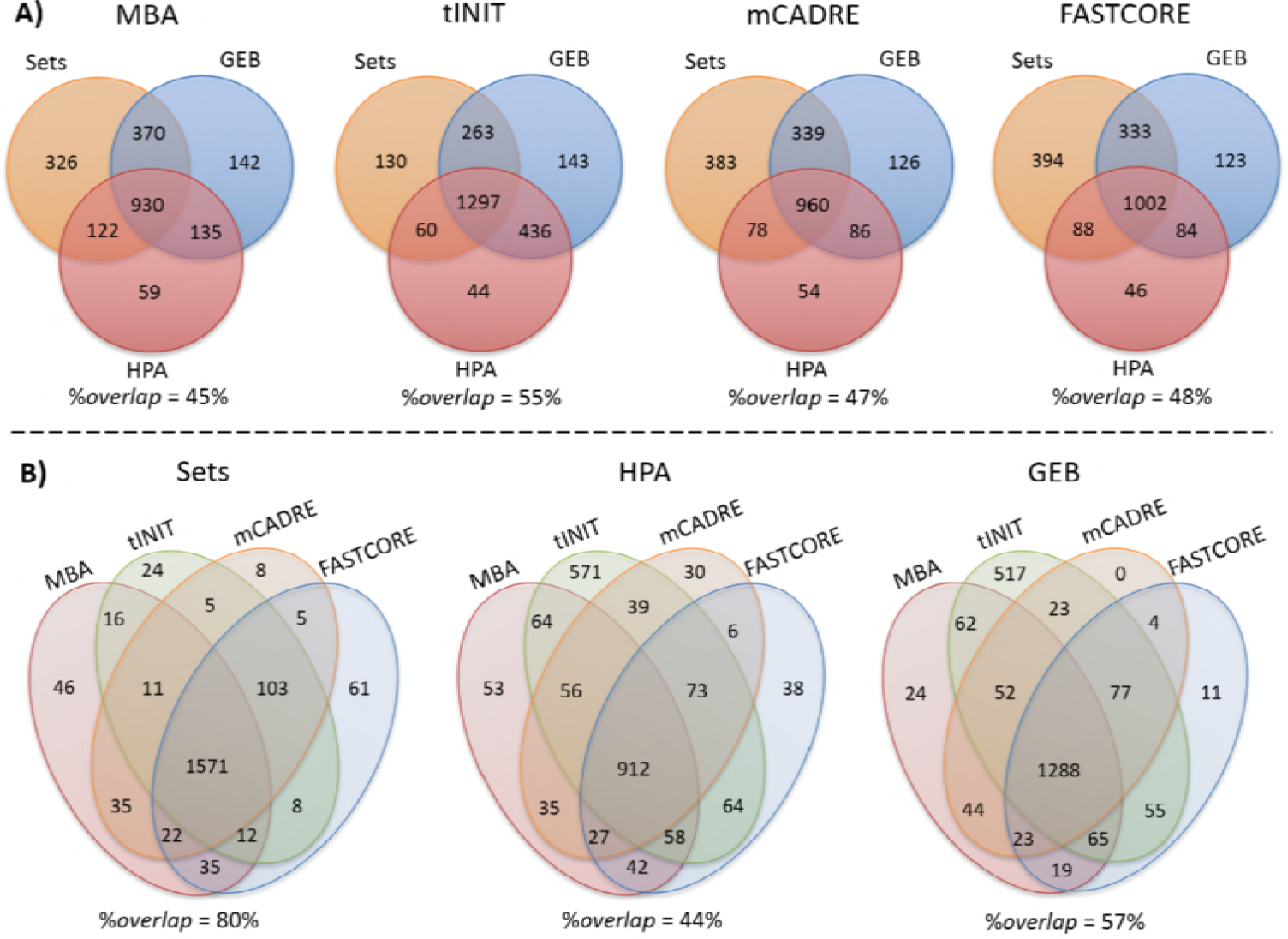
Hepatocytes’ metabolic models reaction overlap considering: (A) the same algorithm; (B) the same omics data source. The overlap percentage was calculated by dividing the number of overlapping reactions by the number of reactions present in the union of the models.

Considering the models generated by the same algorithm, it is clear that MBA has a smaller overlap (only 930 reactions) compared to the other methods. This could be explained by the stochastic nature of the algorithm, which generates different models in each iteration.

To analyze the influence of different algorithms or data sources in the final models, we calculated the percentage of overlapping reactions. These values show that the same input data under different algorithms produces metabolic models with lower variance, i.e with more overlap, than using the same algorithm for different omics data sources. Furthermore, the mean of reactions that belong to all models of the same algorithm is around 49%, and around 60% when the models are grouped by data source. Again, the variability of the final results seems to be dominated by the data source factor. This can also be observed performing a hierarchical clustering of the models (Fig 5 A).

**Fig 5.**
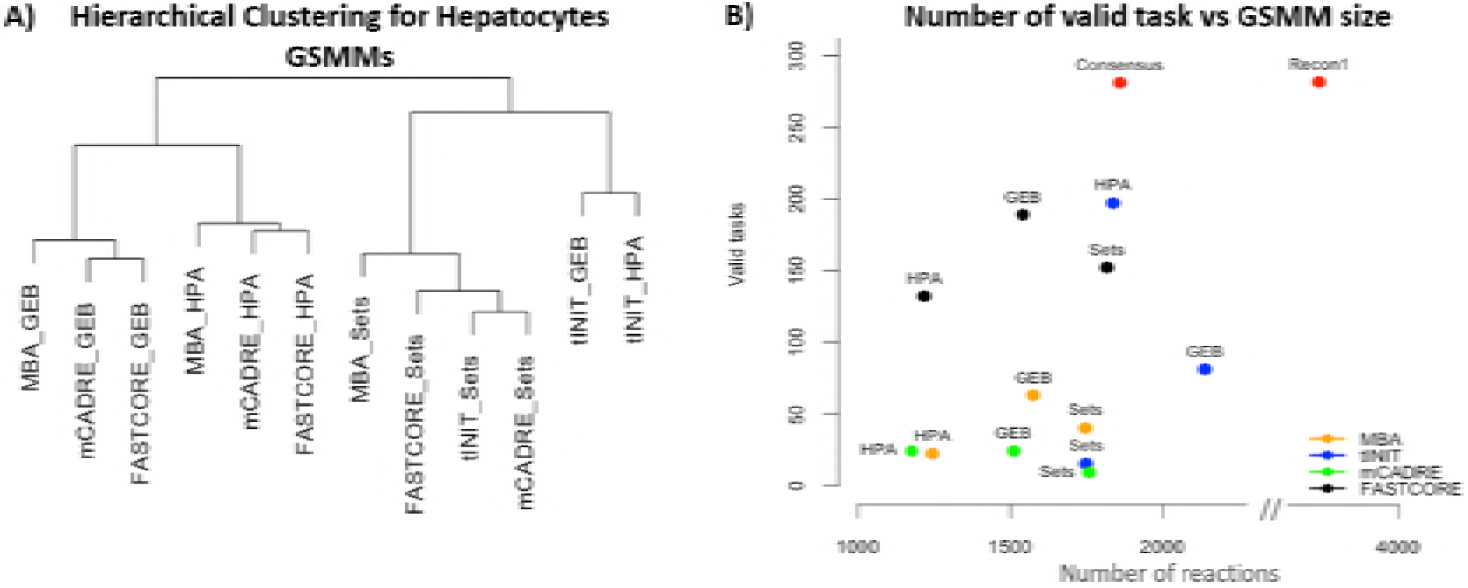
Comparison of the models reconstructed by different algorithms and omics data. A) Hierarchical clustering of the twelve hepatocytes models. B) Correlation between satisfied metabolic tasks and number of reactions for each hepatocytes model.

A set of metabolic tasks known to occur in hepatocytes was previously presented by Gille et al. [35]. Some of these tasks are impossible to satisfy using Recon 1 for the phenotype simulations, because these include metabolites which are not present in the model. Thus, these tasks will not be considered in the validation process. We have also removed disease related tasks, as these do not configure the regular behaviour of hepatocytes.

The Recon 1 template model is able to satisfy 281 of the remaining 363 metabolic functions tested, and therefore this number represents the maximum that may be achieved by any model derived from Recon 1 as the template. This set of 281 metabolic tasks was used to validate each hepatocytes metabolic model, running simulations with it to assess if each task could be satisfied.

The relationship between the model size and the number of satisfied tasks for all reconstructed models is presented in Fig 5 B. FASTCORE is able to produce consistent models independently of the input data. tINIT also has a significant percentage of valid tasks when the data source is HPA. However, generically, the number of satisfied metabolic tasks is very low compared with the performance of the template model *Recon 1*.

Using the consensus algorithm proposed in this work, we reconstructed the final hepatocytes consensus model based on the twelve partial models. The consensus model has 1859 reactions satisfying the whole set of 281 metabolic tasks (Fig 5 B). So, this model is able to perform the same tasks as *Recon 1*, while keeping only about 50% of the reactions.

### Glioblastoma tissue-specific metabolic models

Following a similar pipeline to the one described for hepatocytes, eight models were reconstructed for the glioblastoma U-251 cell line. The set of algorithms was the same, while the data sources were limited to HPA and GEB, since no pre-defined lists of reactions were available in this case. Fig 6 shows the overlap of the resulting models from the different algorithms, when each of the omics data source was considered. Fig 6. Reactions overlap of U-251 metabolic models grouped by data source (HPA and GEB).

**Fig 6.**
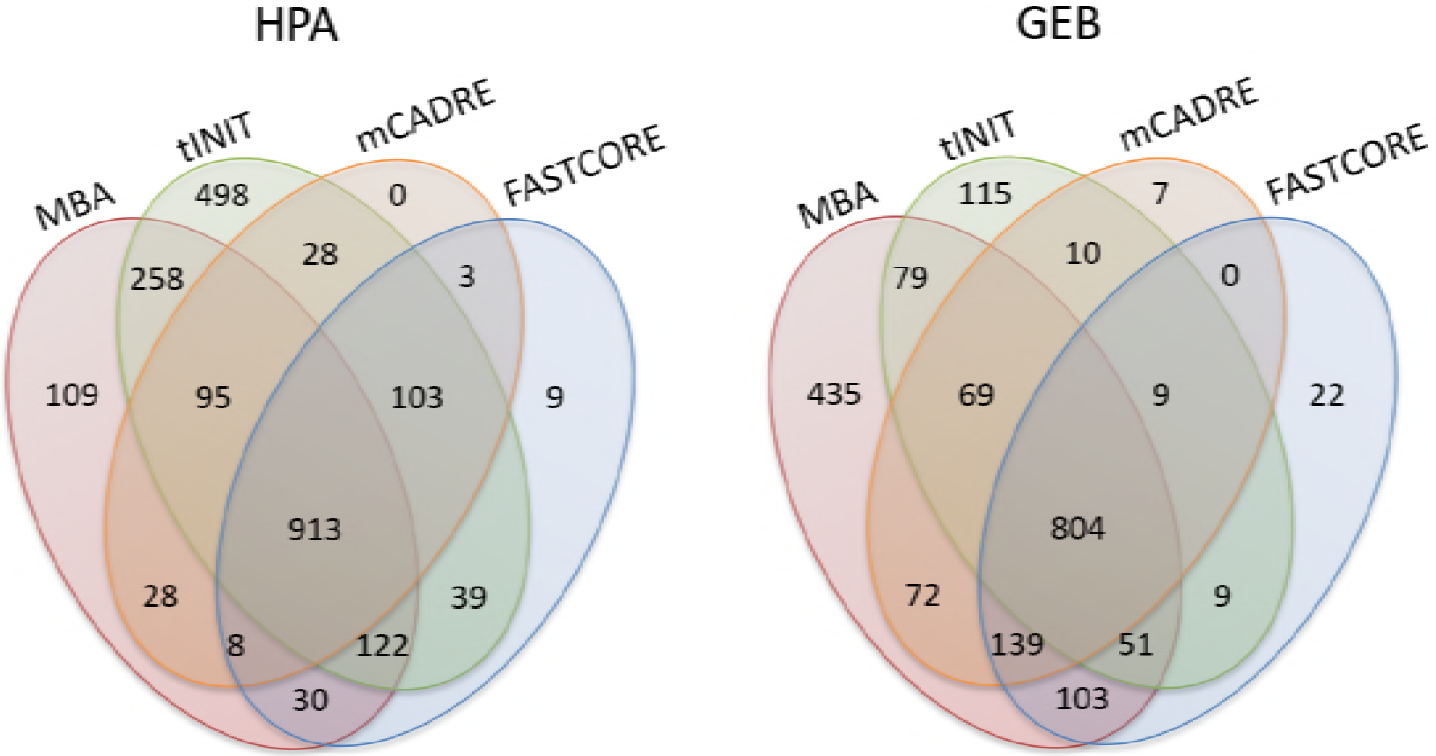
Reactions overlap of U-251 metabolic models grouped by data source (HPA and GEB).

Comparing the models reconstructed with the same data source, tINIT and MBA algorithms produce models with a higher number of exclusive reactions, i.e., reactions present in a single model. The number of reactions shared by all models for each data source is similar, 913 and 804 for HPA and GEB, respectively. However, the intersection of these two sets is only of 577 reactions.

Fig 7 presents the intersection of the same models, but considering the algorithm as the category. MBA has the highest overlap when compared to the other methods, but with the exception of the tINIT_HPA model, the models from MBA have a significant increase in the number of reactions, when compared with the other algorithms. The models from mCADRE and FASTCORE have a similar number of reactions and the ones generated by HPA and GEB are also of a similar size.

**Fig 7.**
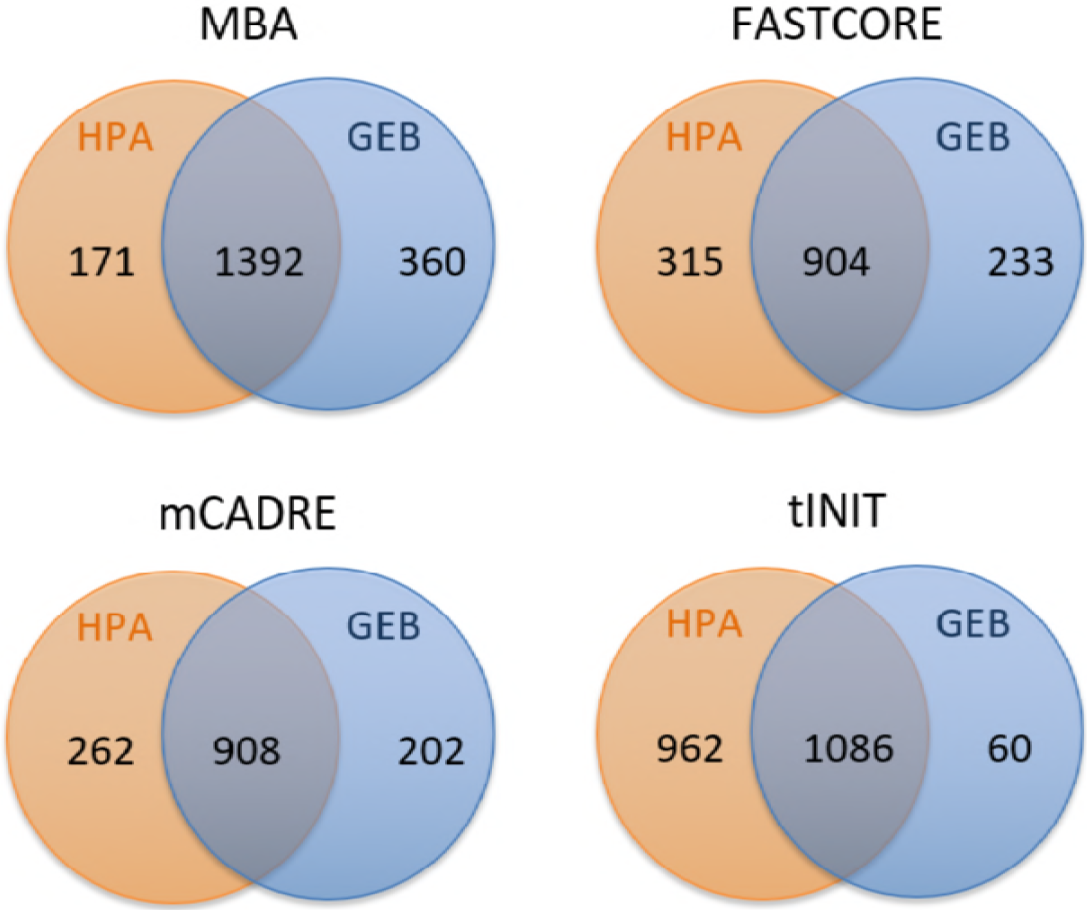
Reactions overlap of U-251 metabolic models grouped by algorithms (MBA, FASTCORE, mCADRE and tINIT).

One of the hallmarks of cancer is the ability that cancer cells have to proliferate. To address this issue, FBA simulations were done to test the growth rate (through maximization of the biomass flux) of each model. The biomass equation was collected from the Recon 2 metabolic model and the RPMI-1640 medium [36] has been considered in all simulations.

As a result, none of the models was able to produce biomass which indicates there are gaps in the models towards the production of some biomass precursor metabolites (essential for growth). We then tested how many biomass precursors could be produced by each metabolic model, by adding artificial drain reactions to excrete each biomass precursor and simulating the maximization of these reactions. Table 3 presents the number of biomass precursors produced by each of the U-251 metabolic models.

**Table 3.**
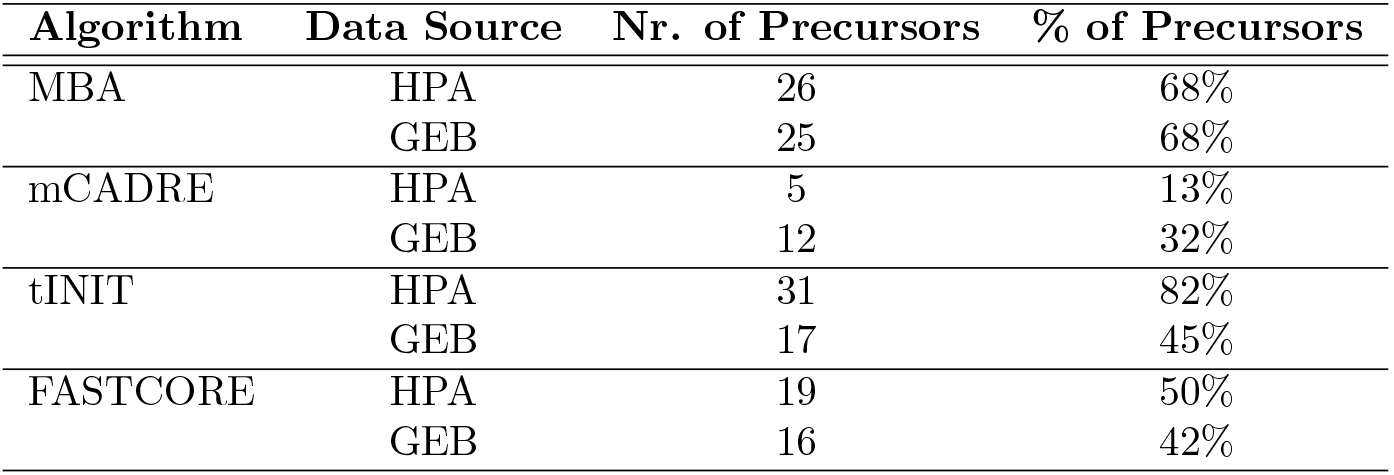
Number of biomass precursors produced by each of the U-251 metabolic models. The biomass equation was obtained from Recon 2 metabolic model which contains 38 precursors metabolites.

The U-251 model generated by the tINIT algorithm using HPA has the highest number of biomass precursors satisfied. This was expected since this model has approximately 500 more reactions than the remaining models.

Given these results, the reconstruction of a single, unified and global U-251 metabolic model capable of sustaining cell growth is desirable. This model must be able to carry flux on the biomass reaction, to allow to simulate the proliferation of cells, predicting their growth rate.

As before, the consensus U-251 metabolic model was built considering all previously reconstructed models by different methods and data sources. The process starts with the reconstruction of the partial models (*pModel*_*i*_ *i* ∈ 1, ..8) as described in the Methods section.

In this case, the tasks will be defined as the production of biomass precursors. So, we checked how many biomass precursors each partial model was able to produce. Fig 8 A) depicts the number of reactions and the number of biomass precursors produced by each *pModel*. The highest decrease on the number of produced biomass precursors occurs between *pModel*_4_ and *pModel*_5_. Thus, the process of reconstructing the final consensus model starts with *pModel*_4_ as the initial model. In each iteration, the algorithm takes as input a partial model (*pModel*_*i*_) and tries to remove the maximum number of reactions that were lost between *pModel*_*i*_ and *pModel*_*i*+1_, maintaining the biomass precursors produced by *pModel*_*i*_ and not by *pModel*_*i*+1_. In this case, the lost reaction set (LRS) and the lost biomass precursors set (LBS) are composed by the difference between the two partial models *pModel*_4_ and *pModel*_5_. At the end of each iteration, the reactions that do not have an influence in the loss of biomass precursors between two partial models (*toDel*_*i*_) are appended to the LRS in the next iteration. The process ends with the processing of *pModel*_0_, in this case the full *Recon 1* model.

**Fig 8.**
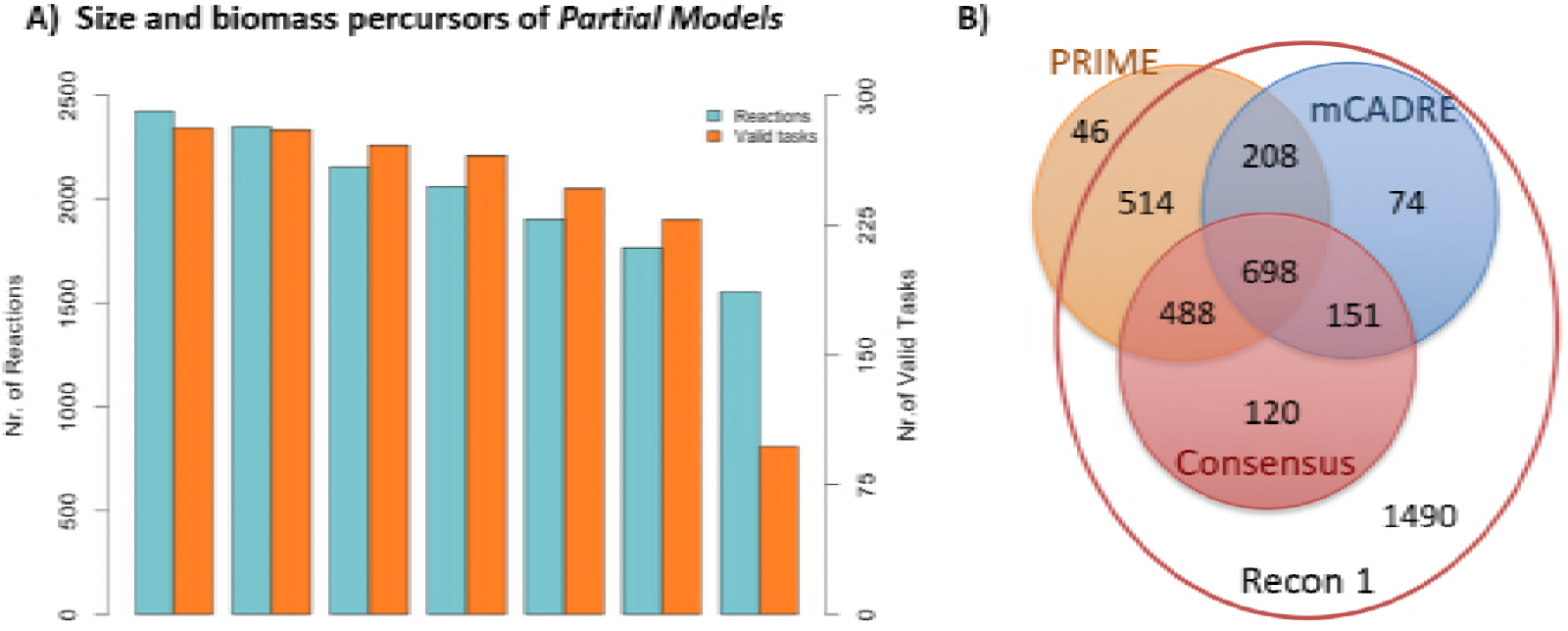
A) Number of reactions (green bars) and number of biomass precursors that can be produced (orange bars) by the *pModels*. The *pModel*_8_ was ignored since *pModel*_7_ does not produce any of the biomass precursors. B) Overlap of metabolic models. The PRIME and mCADRE models are available in the methods publication articles. The consensus model is our model, reconstructed during this study.

The final consensus model obtained is composed of 922 genes, 1376 metabolites and 1457 reactions. This model is able to simulate the biomass production, through FBA, using the RPMI-1640 medium [36]. The flux rate for biomass equation is around 0.0291 *mmol/gDW/hr*. Although the lower biomass flux rate, when compared with the original model Recon 1 (0.084 *mmol/gDW/hr*), this process is able to achieve a final consensus model based in all previous models capable to simulate the biomass production.

### Other tissue-specific metabolic models

Glioblastoma GSMMs were already reconstructed in previous studies [21,23]. The glioblastoma tumor cells and U-251 cell line GSMMs reconstructed by mCADRE and PRIME algorithms were used to perform a comparison with our consensus model. The overlap between these glioblastoma metabolic models is provided in Fig 8 B).

Analyzing the model obtained by PRIME, we verified that the Recon 1 template model used by the algorithm is not the original model, but an extended version which has 46 extra reactions. These reactions are essentially for excretion of cytosol metabolites which can lead to significant differences in the phenotype simulation results. The models PRIME, mCADRE and Consensus are composed by 1952, 1131 and 1457 reactions, respectively.

To further validate our model we decided to perform additional phenotype simulations, using the methods pFBA [5], iMAT [37], GIMME [38] and E-Flux [39], which are able to integrate omics data to improve prediction results. Transcriptomics data published by Gholami et al. [40], which were not used for model reconstruction, were used as inputs. Experimental flux values published by Jain et al. [41] were used to compare with the flux exchange rates predicted by the different methods and the normalized prediction errors were calculated using the equation:

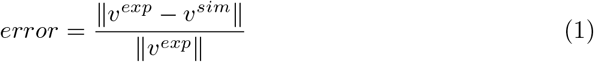

where *ν*^*exp*^ is the vector of measured flux values and *ν*^*sim*^ is the vector of predicted values.

In this study, the glioblastoma metabolic model obtained by the mCADRE reconstruction method was not considered, because this model is not able to grow when simulated using the *Recon2* biomass equation, even with the removal of the metabolites present in the biomass equation and not in the model. The normalized prediction errors are given in Table 4.

**Table 4.** The normalized prediction errors, associated with the simulation methods pFBA, GIMME, E-Flux and iMAT, for U-251 model reconstructed by PRIME and consensus algorithm.

|  | pFBA | E-Flux | GIMME | iMAT |
| --- | --- | --- | --- | --- |
| CONSENSUS | 1.2866 | 0.7427 | 0.7639 | 0.7102 |
| PRIME | 0.7849 | 2.9739 | 0.7481 | 0.7461 |

The best combinations have a normalized error around 0.7-0.8. The one reaching a lower prediction error was obtained with our consensus U-251 metabolic model using the iMAT simulation method. Although no definitive conclusions may be taken from these results, this contributes to provide a higher confidence in our consensus model.

### Critical genes

The validation of metabolic models is a hard task when fluxomics data are not available. So, further tests were done to check if the consensus metabolic model has a better phenotype prediction capability than the global model Recon 1.

As a final test, we calculated the predicted critical genes of both models (our consensus glioblastoma model vs the Recon 1). We considered to be critical (or essential) the genes that inhibit growth when they are removed from the model. We obtained these gene sets through FBA simulations, testing each gene present in the model by performing its knock out, i.e. the reactions associated through GPRs to this gene were constrained to have no flux.

The final consensus model of the U-251 cell line has 89 critical genes, of which 80 are also critical genes in Recon 1. Thus, nine genes are only critical on the U-251 metabolic model - *G6PT2*, *SLC5A7*, *NME2*, *NME1*, *SLC6A14*, *PTDSS1*, *SLC16A10*, *CDS1*, *CTPS*. Remarkably, most of these genes have been associated to cancer cell growth in several research studies. The function and the relevance of these genes on glioblastoma cancer cells are detailed next:

- The **G6PT2** gene regulates the Glucose-6P transport from cytoplasm to the lumen of the endoplasmic reticulum. Studies demonstrate that intracellular signalling and invasive phenotype of brain tumor cells could be regulated by this gene [42]. Moreover, silencing the G6PT gene in U-87 brain tumor-derived glioma cells may induce necrosis and late apoptosis [43]. Thus, control of the G6PT expression can lead to the development of new strategies to prevent cancer development in glial cells.
- The **SLC5A7** gene encodes a high-affinity choline transporter. Choline is used for the synthesis of essential lipid components of cell membranes [44]. A higher choline concentration in the cells has been related with cell proliferation and malignant progression of cancer [45,46] being the abnormal choline metabolism considered, by Glunde et al., as a new hallmark of cancer [47]. Kumar et al. [48] demonstrate that using specific choline kinase inhibitors may be a promising new strategy for the treatment of brain tumors.
- The **SLC6A14** gene encodes the protein called *sodium- and chloride-dependent neutral and basic amino acid transporter B(0+)* which can transport all essential amino acids, as well as glutamine, arginine, and asparagine [49]. Cancer cells, to support their rapid cell growth, induce the over-expression of this gene. This phenomenon has been observed in cervical cancer, colorectal cancer and breast cancer cell lines [50–52]. The SLC6A14 deletion was studied in mouse models of breast cancer by Badu et al. [53]. The study demonstrated that the development and progression of breast cancer were markedly decreased *in vitro* and *in vivo* when SLC6A14 is deleted.
- The **CDS1** is a protein coding gene which regulates the amount of phosphatidylinositol available for signaling by catalyzing the conversion of phosphatidic acid to CDP-diacylglycerol. CDP-diacylglycerol is an important precursor for the synthesis of phosphatidylinositol (PtdIns), phosphatidylglycerol, and cardiolipin [54,55]. The cardiolipin compound is one of the biomass precursors present in Recon 2 biomass equation. Thus, its production is essential.
- The **PTDSS1** gene encodes phosphatidylserine synthase 1 (PSS1) which is involved in the production of phosphatidylserine. This gene is involved in a patent related to the development of a molecular-based method of cancer diagnosis and prognosis. Together with five others genes, the PTDSS1 has a higher expression in tumor samples when compared with control samples [56].
- The **CTPS** gene encodes an enzyme responsible for the conversion of UTP (uridine triphosphate) to CTP (cytidine triphospate). The development of methods and pharmaceutical compositions to inhibit the lymphocyte proliferation through the CPTS1 inhibitors has been protected by a patent [57].
- The **NME2 / NME1** genes were identified as potential tumor suppressors, which reduce the tumor progression and proliferation [58]. Thus, it was unexpected that these genes were essential for the metabolic model. To understand this result, we did a deep analysis of the reactions where these genes are involved. The two genes regulate the activation of nucleoside-diphosphate kinase reactions in the nucleus. These reactions are responsible to produce essential metabolites present in biomass equation, namely Deoxyguanosine triphosphate (dGTP), Deoxycytidine triphosphate (dCTP), Deoxyadenosine triphosphate (dATP) and Deoxythymidine triphosphate (dTTP). These metabolites are used in cells for DNA synthesis.

## Conclusions

In this work, a critical assessment of the most important methods for the reconstruction of tissue-specific metabolic models was performed. Moreover, the consistency of information across important omics data sources was analyzed and these data were used to verify the impact of such differences in the final metabolic models generated by each method. The results show that metabolic models obtained depend more on the data sources used as inputs than on the algorithms used for the reconstruction. To validate the performance of the obtained metabolic models regarding phenotype prediction, a set of metabolic functions was tested for each metabolic model. Generically, the number of satisfied metabolic functions was surprisingly low. This shows that existing methods for the reconstruction of tissue-specific metabolic models, based on a single omics data source, are not enough to generate high quality metabolic models.

Here, a strategy to build a final metabolic model using the combination of generated models through different algorithms and data sources was presented. This process shows that with a similar number of reactions, it is possible to achieve a final model capable of satisfying all possible metabolic tasks.

A variant of the previous method was also used to reconstruct a consensus model of glioblastoma cell lines. The method was able to find a consistent model, able to sustain growth of cancer cells, with around half of the reactions from the template model. The consensus model showed a good predictive ability of flux distributions, combined with further omics data, competitive with previous approaches. Also, it allowed to uncover several candidate essential genes of glioblastoma cell lines, a few of which have been previously identified in literature as relevant targets or biomarkers.

Overall, the method here proposed provides an additional tool in modeling efforts in cancer research and drug development, providing a way to build more robust metabolic models given available omics data sources and phenotypic data.

## Supporting information

**S1 File. Hepatocytes Consensus Model.**

**S2 File. Glioblastoma cell line Consensus Model.**

## Acknowledgments

We would like to thank Paulo Vilaça for discussions during the software development.

